# BUSZ: Compressed BUS files

**DOI:** 10.1101/2022.12.19.521034

**Authors:** Pétur Helgi Einarsson, Páll Melsted

## Abstract

**Summary:** We describe a compression scheme for BUS files and an implementation of the algorithm in the bustools software. Our compression algorithm yields smaller file sizes than gzip, at significantly faster compression and decompression speeds. We evaluated our algorithm on 533 BUS files from scRNA-seq experiments with a total size of 1Tb. Our compression is more than 2x faster than the fastest gzip option and results in 1.5x smaller files than the best gzip compression. This amounts to an 8.3x reduction in the file size, resulting in a compressed size of 122Gb for the dataset.

**Availability and Implementation:** A complete description of the format is available at https://github.com/BUStools/BUSZ-format and an implementation at https://github.com/BUStools/bustools

**Contact:** online.

## 1 Introduction

The Barcode-Umi-Set (BUS) file format (Melsted *et al*., 2019) was designed to represent intermediate results from single-cell RNA-sequencing (scRNA-seq) experiments. The goal was to separate the process of alignment and downstream analysis allowing for rapid analysis and adaptation to different scRNA-seq technologies. The modular design of bustools (Melsted *et al*., 2021) allows for using various sub-commands together to build pipelines tailored for various projects or technologies, which is achieved by streaming or storing intermediate BUS files.

The original BUS files are often smaller than the corresponding FASTQ files containing the original sequences, especially after sorting. The speed of analysis makes it convenient to store or archive the sorted BUS files for reproducibility or future analysis. The BUS format was designed for fast and modular processing, but still has room for compression. A simple method would be to use gzip for compression, but the structured format of the data allows for tailoring the compression method to the needs of bustools.

The first release of the Human Cell Atlas (Consortium* *et al*., 2022) contained over 500,000 cells. Advances in scRNA-seq technologies and increased sequencing throughput have already made it feasible for single labs to generated datasets of several million cells. Just the raw sequencing data can be estimated to be 2Tb per 1 million cells. Thus, to contain the cost of storing and transferring BUS files we propose a compression method to efficiently compress and decompress BUS files and implement it into bustools.

## 2 Methods

Technologies for scRNA-seq experiments vary in how they measure gene expression of isolated cells and capture information about individual molecules. The BUS file abstracts these technological differences by encoding the barcode of each cell and the unique molecular identifier (UMI) as short oligonucleotides encoded as integers. For each molecule the corresponding read from the cDNA is not stored, but rather the transcript or set of transcripts it aligned to is stored as an equivalence class (EC). The EC corresponds to a set of transcripts, each set is encoded as a unique integer and the map is stored alongside of the BUS file.

A BUS file consists of a header followed by a sequence of BUS records (Melsted *et al*., 2019). Each BUS record occupies exactly 32 bytes and consists of six fields; barcode, unique molecular identifier (UMI), equivalence class (EC), read count, flags, and padding. These fields are 8, 8, 4, 4, 4, 4 bytes in size, respectively. The barcodes and UMIs represent nucleotide sequences and are two-bit encoded, allowing them to be expressed as unsigned 64-bit integers. The fixed length format of BUS records allows for fast loading of BUS files and circumventing the need for parsing textual data.

The layout of records in BUS files is row-based, as shown in Figure 1A. To compress the BUS file we use a columnar layout for the compression, where the columns correspond the fields of the BUS records, similar to the CRAM format (Fritz *et al*., 2011).

**Figure 1:**
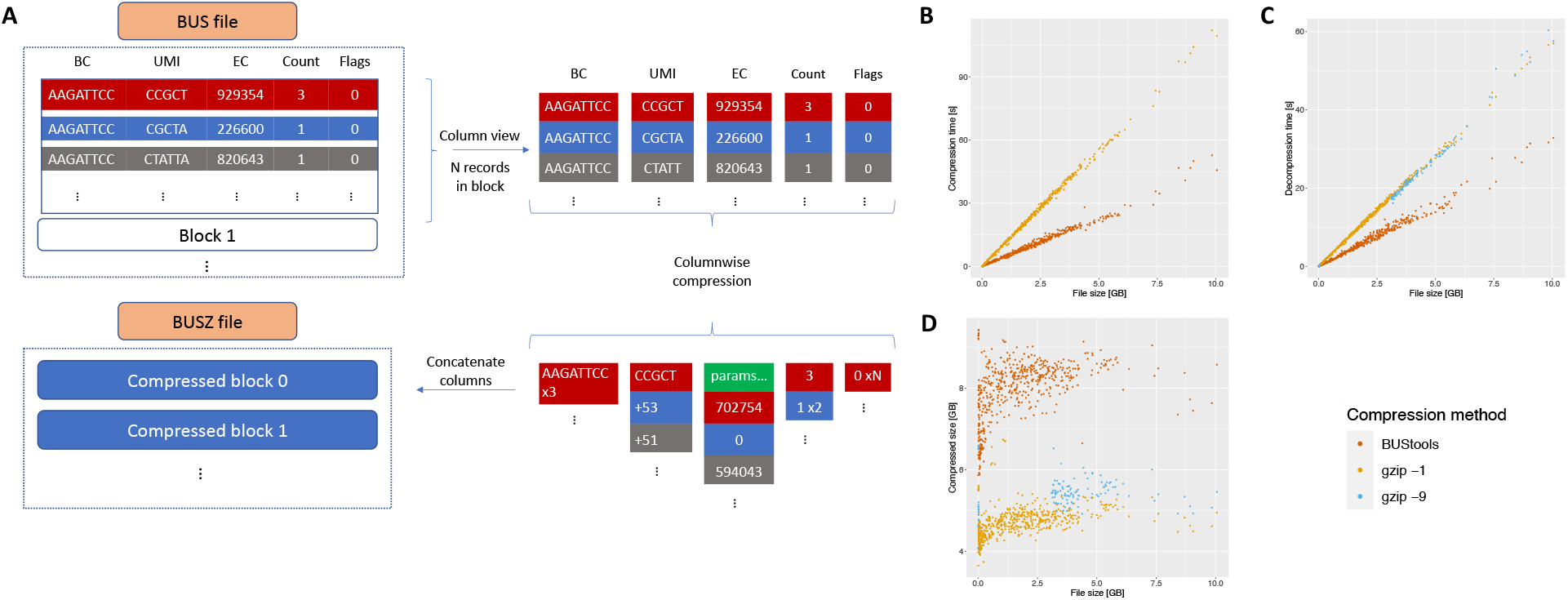
**(A)** A general outline of the compression scheme for BUS files. **(B)** Comparing compression time between BUStools and `gzip −1`. The best (`gzip −9`) option is omitted due to extremely long compression time. **(C)** Comparing decompression time of the three methods. **(D)** Comparing compression ratios using the three methods.

Our compression algorithm assumes a sorted input, which can be obtained using the `bustools sort` command. The input is sorted lexicographically by barcodes first, then by UMIs, and finally by the equivalence classes. This can be seen as the first level of compression, as records with the same barcode, UMI, and EC are merged. We split the input into blocks of *N* records (by default *N* = 100000). Within each block, the columns are compressed independently, each with a customized compression-decompression scheme (codec). The padding column is not included in the compressed file.

The compression scheme of each column is as follows. To simplify the notation, we let *RLE*_0_ denote run-lengthencoding on zeros and *FRLE*_0_ denote *RLE*_0_ followed by Fibonacci encoding (Fraenkel and Klein, 1996). Since Fibonacci encoding does not encode zeros, we increment each value by one. Similarly, we define *RLE*_1_ and *FRLE*_1_ for one instead of zero.

When compressing the barcode column we expect that many BUS records originate from reads captured from the same cell, thus resulting in a repeat of barcodes. We take the differences of adjacent barcodes, which are non-negative since the barcode values are increasing. Since this list is expected to contain a run of zero differences, we encode it with an *FRLE*_0_.

The UMIs are encoded in a similar manner as the barcodes. However, they only increase within the same barcode so we must modify how the differences are computed. The difference is only taken of adjacent records from the same cell. Otherwise, the absolute value is used. We then continue with *FRLE*_0_ as before.

The third column, containing the equivalence classes, is not as well structured as the first two as it has higher entropy and is harder to compress (Supplementary Material). To compress this column we modified a variant of PForDelta (Zukowski *et al*., 2006) called NewPFD (Yan *et al*., 2009). This codec splits the list of ECs into sub-blocks of *N_PFD_* consecutive values (by default *N_PFD_* = 512). For each sub-block ***B*** we find two parameters, *k* and *b*, so that *f* (default *f* = 0.9) fraction of values fall in the interval [*k, k* + 2^*b*^ −1]. We choose *k* to be the smallest value in the sub-block, whereas *b* is computed as the number of bits required to encode *f* · *N_PFD_* of the values (*x - k*), *x* ∈ ***B***. Any value requiring more than *b* bits is considered an exception and the remaining high bits are Fibonacci encoded. A complete description of the encoding is in the Supplementary Materials.

The last two columns, count and flags, are compressed using *RLE* and Fibonacci encoding. The count values are strictly positive and are most often equal to one, so we use run-length encoding on ones (*FRLE*_1_). The flags column contains mostly zeros so we use *FRLE*_0_.

Each compressed block is preceded by a block header, containing the number of records contained in the block, as well as the size in bytes of the compressed block.

The compressed file starts with a header containing a fixed magic number to identify the file as a compressed BUS file, the header values of the input file, and the parameters used for the compression. Optionally, we output an indexing file, which contains the first barcode of each block and the size of the block, facilitating fast lookup of records for specific barcodes without decompressing the entire BUSZ file.

## 3 Results

To evaluate our compression algorithm, we measured the time it took to compress and decompress 533 BUS files collected as a part of a larger survey (Booeshaghi *et al*., 2022). The total size of the dataset is 1010Gb of BUS files. All experiments were performed using a single core on a Intel Xeon CPU E5-2697 2.7GHz processor. File sizes ranged from 4MB to 10GB and were all sorted prior to compression. We compare the performance with gzip, using the fastest compression (`gzip −1`) and the best compression (`gzip −9`). To ensure that disk access patterns and caching would not affect the results, we eliminate disk read latency by caching the input files before running each compression method. Since compressing the files using gzip with the best option (−9) takes substantially more time than the other two methods, we only used that option to compress the 100 smallest and the 100 largest BUS files. The results in Figure 1 were acquired using a block size of *N* = 100000 and a NewPFD sub-block size of *N_PFD_* = 512.

Our method performed consistently better than gzip, both in terms of speed and compression ratio. For compression and decompression, all methods have a running time roughly linear in terms of input size up to 10GiB. Fitting the data with a linear model shows a speedup of 2.4x and 60x compared to gzip with options *-1* and *-9*, respectively (Fig 1A). Similarly, BUStools’ decompression time is 1.9x faster than both options for gzip (Fig 1B). We estimate the compression ratio using a linear model comparing the original and compressed size and find a compression ratio of 8.3, 4.9, and 5.4 for BUStools, `gzip −1`, and `gzip −9`, respectively (Fig 1C).

Together these results show that the compression scheme, as implemented in bustools, is significantly faster and shows better compression than gzip alone.

## Acknowledgements

We thank Ángel Gálvez-Merchán and A. Sina Booeshaghi for help with benchmarking. Atli Fannar Franklín worked on an initial prototype of the software. Lior Pachter provided valuable feedback on the design of the compression and use cases.

## Funding

This work was supported by the Icelandic Research Fund Project grant number 218111-051.

## Notes

### Competing Interest Statement

The authors have declared no competing interest.

