## Supplementary methods for "BUSZ: Compressed BUS files"

### Compressing BUS files - Supplement

December 19, 2022

### 1 Fibonacci encoding

As Fibonacci encoding does not guarantee a byte-aligned encoding, we pad a Fibonacci-encoded block with 0-bits such that the number of bytes for that block is a multiple of 4 or 8, depending on the type of integer used for encoding (`uint32_t` or `uint64_t`).

### 2 Compressing the Equivalence Class column

#### 2.1 NewPFD

To compress the EC column, we use a compression scheme (*codec*) called NewPFD. This codec is based on another codec called PFOR Delta (PFD), which in turn is a variant of Patched Frame of Reference (PFOR) with delta encoding, i.e. differences. Since the ECs are not sorted, we skip computing the differences.

The PFD codec compresses lists of integers by splitting the input into chunks of e.g. 128 numbers, and has parameters $b$, and $k$, such that a certain fraction of numbers, $p$, fit in the interval $[k, k + 2^b - 2]$. Any number not in the interval is considered an exception. Let $B$ be a block of numbers to encode. To encode $B$, we allocate $b * |B|$ bits as a primary encoding block; $B_p$. Each number $x \in B$ is encoded in $B_p$ using exactly $b$ bits as if it fits the interval defined by $b$ and $k$. Exceptions are indicated with $b$ 1-bits in $B_p$ and their values are encoded after $B_p$ using any codec, e.g. Fibonacci encoding or Elias Delta encoding.

In NewPFD, we store the lower $b$ bits of the exceptions in $B_p$. The exceptions are then stored in two additional vectors. The former vector stores the gaps between the indices of exceptions in $B_p$, whereas the latter stores the overflowing high bits. These vectors can then be compressed by any codec.

Our implementation of NewPFD encodes the parameters, number of exceptions, along with the two exception vectors using Fibonacci encoding prior to the encoding of $B_p$. This is due to the padding we described in Section 1. For the Fibonacci encoding, we use `uint32_t` to save bits lost to padding.

#### 2.2 Other methods

We tested several other methods to compress the ECs. These methods were composed of zero or more list encoders, followed by an integer encoder. We call a compression scheme with multiple layers of encoders a stack. For each of the following stacks, we tested compression with one of several integer encoder.

- *empty*
- XOR
- XOR - RLE
- XOR - differences
- XOR - differences - RLE

The following integer encoders were tested:

- Elias Delta
- Elias Delta Rice, $k \in \{2, 4, 8, 16, 24, 32\}$
- Exponential Golomb, $k \in \{2, 3, 5, 8, 13, 21\}$

- Exponential Elias, $k \in \{2, 3, 5, 8, 13, 21\}$
- Fibonacci encoding
- Huffman encoding

Exponential Elias uses the same decomposition of an integer into a bucket and an offset as Exponential Golomb, but encodes them using Elias Delta encoding, instead of unary encoding and fixed-width binary encoding.

In addition to the stacks created by the cross product of the above lists, we tested NewPFD using sub-block sizes 128, 256, 512, 1024, and 2048.

| method | 0.bus | 1.bus | 2.bus | 3.bus | 4.bus | 5.bus | 6.bus |
|---|---|---|---|---|---|---|---|
| Increment - EliasDelta | 1.462494 | 1.199965 | 1.199594 | 1.463055 | 1.462947 | 1.256509 | 1.256486 |
| Xor - RunLength - Increment - EliasDelta | 1.474521 | 1.272926 | 1.275875 | 1.474656 | 1.474424 | 1.260942 | 1.261043 |
| Xor - Increment - EliasDelta | 1.475165 | 1.273768 | 1.276745 | 1.475315 | 1.475070 | 1.261339 | 1.261443 |
| Xor - Delta - NegToPosMap - RunLength - Increment - EliasDelta | 1.443001 | 1.202311 | 1.202463 | 1.443144 | 1.443006 | 1.239592 | 1.239594 |
| Xor - Delta - NegToPosMap - Increment - EliasDelta | 1.443566 | 1.202651 | 1.202808 | 1.443722 | 1.443570 | 1.239937 | 1.239943 |
| Increment - EliasRiceDelta_2 | 1.534855 | 1.207108 | 1.206685 | 1.535179 | 1.535124 | 1.283847 | 1.283807 |
| Xor - RunLength - Increment - EliasRiceDelta_2 | 1.544424 | 1.285990 | 1.289085 | 1.544474 | 1.544294 | 1.281015 | 1.281107 |
| Xor - Increment - EliasRiceDelta_2 | 1.546074 | 1.288250 | 1.291434 | 1.546172 | 1.545952 | 1.282062 | 1.282159 |
| Xor - Delta - NegToPosMap - RunLength - Increment - EliasRiceDelta_2 | 1.518406 | 1.211566 | 1.211682 | 1.518475 | 1.518344 | 1.257091 | 1.257084 |
| Xor - Delta - NegToPosMap - Increment - EliasRiceDelta_2 | 1.520156 | 1.212445 | 1.212572 | 1.520270 | 1.520103 | 1.258130 | 1.258129 |
| Increment - EliasRiceDelta_4 | 1.540622 | 1.261038 | 1.260409 | 1.540954 | 1.540918 | 1.355031 | 1.355002 |
| Xor - RunLength - Increment - EliasRiceDelta_4 | 1.547088 | 1.322292 | 1.325159 | 1.547016 | 1.546903 | 1.355494 | 1.355585 |
| Xor - Increment - EliasRiceDelta_4 | 1.549840 | 1.326270 | 1.329291 | 1.549845 | 1.549662 | 1.357448 | 1.357551 |
| Xor - Delta - NegToPosMap - RunLength - Increment - EliasRiceDelta_4 | 1.520577 | 1.237942 | 1.237895 | 1.520538 | 1.520462 | 1.305612 | 1.305578 |
| Xor - Delta - NegToPosMap - Increment - EliasRiceDelta_4 | 1.523494 | 1.239467 | 1.239442 | 1.523541 | 1.523398 | 1.307480 | 1.307458 |
| Increment - EliasRiceDelta_8 | 1.615182 | 1.297321 | 1.296886 | 1.615861 | 1.615757 | 1.364360 | 1.364330 |
| Xor - RunLength - Increment - EliasRiceDelta_8 | 1.612999 | 1.361168 | 1.363790 | 1.612803 | 1.612793 | 1.361516 | 1.361598 |
| Xor - Increment - EliasRiceDelta_8 | 1.618382 | 1.368776 | 1.371689 | 1.618335 | 1.618206 | 1.365071 | 1.365173 |
| Xor - Delta - NegToPosMap - RunLength - Increment - EliasRiceDelta_8 | 1.576089 | 1.293037 | 1.293136 | 1.575898 | 1.575973 | 1.336855 | 1.336825 |
| Xor - Delta - NegToPosMap - Increment - EliasRiceDelta_8 | 1.581740 | 1.296038 | 1.296178 | 1.581719 | 1.581672 | 1.340387 | 1.340377 |
| Increment - EliasRiceDelta_16 | 1.882349 | 1.465610 | 1.464820 | 1.882349 | 1.882349 | 1.556078 | 1.555942 |
| Xor - RunLength - Increment - EliasRiceDelta_16 | 1.868609 | 1.473411 | 1.474260 | 1.868227 | 1.868537 | 1.562963 | 1.562935 |
| Xor - Increment - EliasRiceDelta_16 | 1.882323 | 1.490345 | 1.491800 | 1.882317 | 1.882319 | 1.571836 | 1.571861 |
| Xor - Delta - NegToPosMap - RunLength - Increment - EliasRiceDelta_16 | 1.787731 | 1.416591 | 1.415978 | 1.787317 | 1.787614 | 1.513006 | 1.512872 |
| Xor - Delta - NegToPosMap - Increment - EliasRiceDelta_16 | 1.801522 | 1.423411 | 1.422885 | 1.801520 | 1.801512 | 1.521577 | 1.521492 |
| Increment - EliasRiceDelta_24 | 1.279998 | 1.280000 | 1.280000 | 1.279998 | 1.279998 | 1.280000 | 1.279999 |
| Xor - RunLength - Increment - EliasRiceDelta_24 | 1.270664 | 1.261671 | 1.261055 | 1.270405 | 1.270617 | 1.271340 | 1.271292 |
| Xor - Increment - EliasRiceDelta_24 | 1.279998 | 1.280000 | 1.280000 | 1.279998 | 1.279998 | 1.280000 | 1.279999 |
| Xor - Delta - NegToPosMap - RunLength - Increment - EliasRiceDelta_24 | 1.269751 | 1.271888 | 1.271780 | 1.269446 | 1.269673 | 1.271077 | 1.271027 |
| Xor - Delta - NegToPosMap - Increment - EliasRiceDelta_24 | 1.279998 | 1.280000 | 1.280000 | 1.279998 | 1.279998 | 1.280000 | 1.279999 |
| Increment - EliasRiceDelta_32 | 0.969696 | 0.969697 | 0.969697 | 0.969696 | 0.969696 | 0.969697 | 0.969697 |
| Xor - RunLength - Increment - EliasRiceDelta_32 | 0.962626 | 0.955814 | 0.955347 | 0.962431 | 0.962591 | 0.963139 | 0.963103 |
| Xor - Increment - EliasRiceDelta_32 | 0.969696 | 0.969697 | 0.969697 | 0.969696 | 0.969696 | 0.969697 | 0.969697 |
| Xor - Delta - NegToPosMap - RunLength - Increment - EliasRiceDelta_32 | 0.961935 | 0.963554 | 0.963472 | 0.961704 | 0.961876 | 0.962940 | 0.962902 |
| Xor - Delta - NegToPosMap - Increment - EliasRiceDelta_32 | 0.969696 | 0.969697 | 0.969697 | 0.969696 | 0.969696 | 0.969697 | 0.969697 |
| Increment - ExpGolomb_2_Unary_Binary | 1.197293 | 0.928346 | 0.927945 | 1.197700 | 1.197625 | 0.991623 | 0.991589 |
| Xor - RunLength - Increment - ExpGolomb_2_Unary_Binary | 1.203037 | 0.990634 | 0.993249 | 1.203041 | 1.202902 | 0.991273 | 0.991354 |
| Xor - Increment - ExpGolomb_2_Unary_Binary | 1.204034 | 0.991972 | 0.994640 | 1.204067 | 1.203906 | 0.991901 | 0.991985 |
| Xor - Delta - NegToPosMap - RunLength - Increment - ExpGolomb_2_Unary_Binary | 1.172255 | 0.921903 | 0.921950 | 1.172277 | 1.172185 | 0.966512 | 0.966502 |
| Xor - Delta - NegToPosMap - Increment - ExpGolomb_2_Unary_Binary | 1.173296 | 0.922411 | 0.922465 | 1.173347 | 1.173232 | 0.967127 | 0.967120 |
| Increment - ExpGolomb_3_Unary_Binary | 1.243830 | 0.956080 | 0.955655 | 1.244268 | 1.244191 | 1.023299 | 1.023264 |
| Xor - RunLength - Increment - ExpGolomb_3_Unary_Binary | 1.248588 | 1.019597 | 1.022269 | 1.248565 | 1.248440 | 1.022171 | 1.022253 |
| Xor - Increment - ExpGolomb_3_Unary_Binary | 1.250017 | 1.021487 | 1.024234 | 1.250034 | 1.249877 | 1.023060 | 1.023147 |
| Xor - Delta - NegToPosMap - RunLength - Increment - ExpGolomb_3_Unary_Binary | 1.215478 | 0.948435 | 0.948472 | 1.215469 | 1.215395 | 0.995917 | 0.995905 |
| Xor - Delta - NegToPosMap - Increment - ExpGolomb_3_Unary_Binary | 1.216970 | 0.949152 | 0.949198 | 1.217006 | 1.216898 | 0.996788 | 0.996779 |
| Increment - ExpGolomb_5_Unary_Binary | 1.348564 | 1.016787 | 1.016307 | 1.349080 | 1.348987 | 1.093077 | 1.093038 |
| Xor - RunLength - Increment - ExpGolomb_5_Unary_Binary | 1.350213 | 1.082010 | 1.084757 | 1.350113 | 1.350036 | 1.089977 | 1.090057 |
| Xor - Increment - ExpGolomb_5_Unary_Binary | 1.352725 | 1.085205 | 1.088079 | 1.352696 | 1.352558 | 1.091494 | 1.091583 |
| Xor - Delta - NegToPosMap - RunLength - Increment - ExpGolomb_5_Unary_Binary | 1.311773 | 1.005987 | 1.005994 | 1.311688 | 1.311661 | 1.060363 | 1.060338 |
| Xor - Delta - NegToPosMap - Increment - ExpGolomb_5_Unary_Binary | 1.314382 | 1.007197 | 1.007220 | 1.314374 | 1.314284 | 1.061843 | 1.061827 |
| Increment - ExpGolomb_8_Unary_Binary | 1.540724 | 1.123583 | 1.122999 | 1.541408 | 1.541274 | 1.217125 | 1.217075 |
| Xor - RunLength - Increment - ExpGolomb_8_Unary_Binary | 1.532470 | 1.187460 | 1.190155 | 1.532275 | 1.532286 | 1.209560 | 1.209636 |
| Xor - Increment - ExpGolomb_8_Unary_Binary | 1.537330 | 1.193246 | 1.196166 | 1.537266 | 1.537169 | 1.212366 | 1.212458 |
| Xor - Delta - NegToPosMap - RunLength - Increment - ExpGolomb_8_Unary_Binary | 1.484413 | 1.105177 | 1.105084 | 1.484196 | 1.484268 | 1.173705 | 1.173660 |
| Xor - Delta - NegToPosMap - Increment - ExpGolomb_8_Unary_Binary | 1.489426 | 1.107369 | 1.107305 | 1.489358 | 1.489317 | 1.176427 | 1.176397 |
| Increment - ExpGolomb_13_Unary_Binary | 1.880204 | 1.352838 | 1.352079 | 1.880660 | 1.880639 | 1.481518 | 1.481452 |
| Xor - RunLength - Increment - ExpGolomb_13_Unary_Binary | 1.853652 | 1.396439 | 1.398302 | 1.853369 | 1.853660 | 1.460593 | 1.460612 |
| Xor - Increment - ExpGolomb_13_Unary_Binary | 1.864746 | 1.408934 | 1.411263 | 1.864773 | 1.864805 | 1.466969 | 1.467021 |
| Xor - Delta - NegToPosMap - RunLength - Increment - ExpGolomb_13_Unary_Binary | 1.799803 | 1.310418 | 1.309919 | 1.799533 | 1.799788 | 1.412478 | 1.412373 |
| Xor - Delta - NegToPosMap - Increment - ExpGolomb_13_Unary_Binary | 1.811306 | 1.315219 | 1.314781 | 1.811376 | 1.811373 | 1.418621 | 1.418548 |
| Increment - ExpGolomb_21_Unary_Binary | 1.454543 | 1.454545 | 1.454545 | 1.454543 | 1.454543 | 1.454545 | 1.454545 |
| Xor - RunLength - Increment - ExpGolomb_21_Unary_Binary | 1.443936 | 1.433715 | 1.433015 | 1.443639 | 1.443878 | 1.444702 | 1.444649 |
| Xor - Increment - ExpGolomb_21_Unary_Binary | 1.454543 | 1.454545 | 1.454545 | 1.454543 | 1.454543 | 1.454545 | 1.454545 |
| Xor - Delta - NegToPosMap - RunLength - Increment - ExpGolomb_21_Unary_Binary | 1.442898 | 1.445255 | 1.445132 | 1.442549 | 1.442807 | 1.444403 | 1.444346 |
| Xor - Delta - NegToPosMap - Increment - ExpGolomb_21_Unary_Binary | 1.454543 | 1.454457 | 1.454456 | 1.454543 | 1.454543 | 1.454545 | 1.454545 |
| Increment - ExpElias_2 | 1.190902 | 0.957325 | 0.956856 | 1.191157 | 1.191243 | 1.031649 | 1.031594 |
| Xor - RunLength - Increment - ExpElias_2 | 1.204842 | 1.025532 | 1.028020 | 1.204957 | 1.204794 | 1.014351 | 1.014446 |
| Xor - Increment - ExpElias_2 | 1.205601 | 1.026555 | 1.029081 | 1.205739 | 1.205562 | 1.014826 | 1.014924 |
| Xor - Delta - NegToPosMap - RunLength - Increment - ExpElias_2 | 1.191523 | 0.974050 | 0.974148 | 1.191744 | 1.191582 | 1.006945 | 1.006947 |
| Xor - Delta - NegToPosMap - Increment - ExpElias_2 | 1.192269 | 0.974462 | 0.974566 | 1.192507 | 1.192329 | 1.007395 | 1.007401 |
| Increment - ExpElias_3 | 1.191976 | 0.963593 | 0.963085 | 1.192238 | 1.192325 | 1.043228 | 1.043165 |
| Xor - RunLength - Increment - ExpElias_3 | 1.206444 | 1.030668 | 1.033176 | 1.206524 | 1.206377 | 1.023309 | 1.023402 |
| Xor - Increment - ExpElias_3 | 1.207207 | 1.031702 | 1.034248 | 1.207309 | 1.207146 | 1.023792 | 1.023888 |
| Xor - Delta - NegToPosMap - RunLength - Increment - ExpElias_3 | 1.192914 | 0.977558 | 0.977640 | 1.193119 | 1.192961 | 1.013545 | 1.013542 |
| Xor - Delta - NegToPosMap - Increment - ExpElias_3 | 1.193657 | 0.977973 | 0.978061 | 1.193883 | 1.193710 | 1.014002 | 1.014001 |
| Increment - ExpElias_5 | 1.203649 | 0.984630 | 0.984132 | 1.203904 | 1.203994 | 1.054571 | 1.054517 |
| Xor - RunLength - Increment - ExpElias_5 | 1.215068 | 1.051968 | 1.054492 | 1.215073 | 1.214920 | 1.038661 | 1.038767 |
| Xor - Increment - ExpElias_5 | 1.215844 | 1.053045 | 1.055608 | 1.215868 | 1.215700 | 1.039159 | 1.039269 |
| Xor - Delta - NegToPosMap - RunLength - Increment - ExpElias_5 | 1.200356 | 0.991704 | 0.991725 | 1.200512 | 1.200361 | 1.031486 | 1.031493 |
| Xor - Delta - NegToPosMap - Increment - ExpElias_5 | 1.201109 | 0.992131 | 0.992158 | 1.201286 | 1.201121 | 1.031959 | 1.031968 |
| Increment - ExpElias_8 | 1.259366 | 0.986453 | 0.985948 | 1.259902 | 1.259939 | 1.058783 | 1.058732 |
| Xor - RunLength - Increment - ExpElias_8 | 1.276166 | 1.057358 | 1.059939 | 1.276224 | 1.276056 | 1.045847 | 1.045957 |
| Xor - Increment - ExpElias_8 | 1.277021 | 1.058447 | 1.061067 | 1.277099 | 1.276916 | 1.046351 | 1.046464 |
| Xor - Delta - NegToPosMap - RunLength - Increment - ExpElias_8 | 1.249847 | 1.003533 | 1.003626 | 1.250030 | 1.249911 | 1.037890 | 1.037884 |
| Xor - Delta - NegToPosMap - Increment - ExpElias_8 | 1.250658 | 1.003971 | 1.004070 | 1.250870 | 1.250733 | 1.038369 | 1.038365 |
| Increment - ExpElias_13 | 1.400850 | 1.086642 | 1.086100 | 1.401229 | 1.401159 | 1.157176 | 1.157102 |
| Xor - RunLength - Increment - ExpElias_13 | 1.406588 | 1.161446 | 1.164157 | 1.406895 | 1.406659 | 1.154426 | 1.154539 |
| Xor - Increment - ExpElias_13 | 1.407629 | 1.162758 | 1.165518 | 1.407964 | 1.407708 | 1.155041 | 1.155158 |
| Xor - Delta - NegToPosMap - RunLength - Increment - ExpElias_13 | 1.366659 | 1.070492 | 1.070261 | 1.366697 | 1.366598 | 1.140178 | 1.140173 |
| Xor - Delta - NegToPosMap - Increment - ExpElias_13 | 1.367636 | 1.070989 | 1.070764 | 1.367700 | 1.367581 | 1.140757 | 1.140754 |
| Increment - ExpElias_21 | 1.398575 | 1.156594 | 1.156249 | 1.399089 | 1.398989 | 1.209035 | 1.209014 |
| Xor - RunLength - Increment - ExpElias_21 | 1.409114 | 1.223548 | 1.226249 | 1.409227 | 1.409025 | 1.212827 | 1.212918 |
| Xor - Increment - ExpElias_21 | 1.410157 | 1.225006 | 1.227759 | 1.410295 | 1.410071 | 1.213506 | 1.213603 |
| Xor - Delta - NegToPosMap - RunLength - Increment - ExpElias_21 | 1.380259 | 1.158505 | 1.158643 | 1.380375 | 1.380256 | 1.193052 | 1.193053 |

| | | | | | | | |
|---|---|---|---|---|---|---|---|
| Xor - Delta - NegToPosMap - Increment - ExpElias_21 | 1.381255 | 1.159087 | 1.159233 | 1.381399 | 1.381259 | 1.193684 | 1.193690 |
| Increment - Fibonacci | 1.463494 | 1.157571 | 1.157027 | 1.464035 | 1.463999 | 1.237922 | 1.237883 |
| Xor - RunLength - Increment - Fibonacci | 1.466258 | 1.235161 | 1.238230 | 1.466381 | 1.466208 | 1.234563 | 1.234663 |
| Xor - Increment - Fibonacci | 1.467303 | 1.236586 | 1.239709 | 1.467456 | 1.467253 | 1.235230 | 1.235335 |
| Xor - Delta - NegToPosMap - RunLength - Increment - Fibonacci | 1.437816 | 1.159045 | 1.159105 | 1.437943 | 1.437808 | 1.210198 | 1.210186 |
| Xor - Delta - NegToPosMap - Increment - Fibonacci | 1.438875 | 1.159598 | 1.159666 | 1.439035 | 1.438874 | 1.210843 | 1.210835 |
| NewPFD_128_Fibonacci_Fibonacci | 1.967217 | 1.577698 | 1.577664 | 1.967100 | 1.967206 | 1.658945 | 1.659021 |
| NewPFD_256_Fibonacci_Fibonacci | 1.982457 | 1.587969 | 1.587967 | 1.982003 | 1.982456 | 1.669278 | 1.669246 |
| NewPFD_512_Fibonacci_Fibonacci | 1.984821 | 1.588714 | 1.588697 | 1.984185 | 1.985466 | 1.670386 | 1.670371 |
| NewPFD_1024_Fibonacci_Fibonacci | 1.988689 | 1.591009 | 1.591050 | 1.986672 | 1.987994 | 1.673046 | 1.673299 |
| NewPFD_2048_Fibonacci_Fibonacci | 1.990630 | 1.592162 | 1.592182 | 1.988610 | 1.987252 | 1.674568 | 1.674817 |
| raw_size | 1.000000 | 1.000000 | 1.000000 | 1.000000 | 1.000000 | 1.000000 | 1.000000 |
| gz1 | 1.799837 | 1.677109 | 1.684655 | 1.804122 | 1.802159 | 1.682700 | 1.683474 |
| gz9 | 1.938520 | 1.789956 | 1.802806 | 1.942817 | 1.940772 | 1.753917 | 1.754799 |
